# HEIMDALL: Disentangling tokenizer design for robust transfer in single-cell foundation models

**DOI:** 10.1101/2025.11.09.687403

**Authors:** Ellie Haber, Shahul Alam, Nicholas Ho, Renming Liu, Evan Trop, Shaoheng Liang, Muyu Yang, Spencer Krieger, Jian Ma

## Abstract

Foundation models for single-cell RNA-sequencing (scRNA-seq) data are emerging as powerful tools for single-cell analysis, yet their performance depends critically on how cells are tokenized into model inputs. Single-cell data lack a canonical tokenization scheme, and many design choices in current single-cell foundation models (scFMs) remain heuristic, entangled, and difficult to evaluate. Here, we introduce Heimdall, a unified framework for dissecting and redesigning tokenizers in scFMs. By decomposing existing tokenization strategies into individual design choices, Heimdall enables attribution of the components that underlie robust generalization, allowing more principled design of improved tokenizers. Combining Heimdall with a minimal transformer backbone, we find that tokenizer design is instrumental for generalization in challenging distribution-shift settings such as cross-tissue, cross-species, and cross-gene-panel cell type classification, as well as reverse perturbation prediction. We show that, while tokenizer choice has little effect in scenarios with matched train and test data, it becomes imperative under distribution shift. Rather than identifying a single globally optimal tokenizer, Heimdall reveals that robust transfer depends on a small number of tokenization design axes – especially gene identity, expression encoding, and ordering – that expose different biological priors to the model. In this sense, universal transferability in scFMs still depends on a non-universal tokenizer interface. Together, these findings establish tokenization as a critical design axis in scFMs and provide design principles and reusable infrastructure for more robust scFMs.

## Introduction

Foundation models are transforming biology, enabling breakthroughs in protein structure prediction [1], genomics [2], and clinical medicine [3]. In single-cell biology, a new class of models – single-cell foundation models (scFMs) [4–10] – has attracted growing attention. Pretrained on millions of single-cell transcriptomic profiles from large-scale atlases [11–13] and finetuned for tasks such as cell type annotation, perturbation prediction, and gene regulatory network inference [14], scFMs promise to serve as general-purpose tools for biomedical research. Yet in practice, their performance is inconsistent: bespoke methods tailored to specific tasks often outperform them [15–17], and zero-shot evaluations have revealed various limitations [18].

One underexplored source of these limitations is tokenization: how a cell’s gene expression profile is converted into model inputs. Because a cell is fundamentally an unordered set of genes with continuous expression values, single-cell data lack an established tokenization standard comparable to those in text or images, and existing scFMs adopt different schemes that encode different biological assumptions [19]. This matters because tokenization determines what information the model sees and how it is structured, yet it has received far less scrutiny than other design choices like architecture, model scale, or pretraining data. Without a principled way to isolate tokenizer effects, it is often unclear whether a model’s successes and failures reflect its architecture, its training data, or its input representation. More fundamentally, scFMs may aspire to universal transferability, but their robustness can only be as universal as the tokenizer interface through which biological information is exposed to the model.

Prior benchmarks compare fully pretrained scFMs, measuring performance after finetuning or in zero-shot settings [14, 15, 17, 18]. These comparisons are useful but limited: because models differ in size, training data, architecture, and tokenization all at once, it is hard to attribute performance differences to any single cause. As a result, the field lacks clear guidance on which tokenization design axes matter most under realistic deployment shifts. This gap has practical consequences. A user applying an scFM on a new tissue, a new species, or a spatial transcriptomics platform with a limited gene panel has no principled basis for choosing between models, or for anticipating where a model might fail.

To address this, we introduce Heimdall, a modular framework for systematically evaluating and designing tokenization strategies in scFMs. Heimdall decomposes existing tokenizers into three functional components – gene identity encoding, expression encoding, and cell construction – and reimplements tokenizers from five leading scFMs within this common framework. This allows us to hold all other factors constant and measure the contribution of tokenization alone, enabling comparisons that prior benchmarks could not make. Using Heimdall, we show that tokenization has little effect when models are tested in the same setting they were trained in, but becomes a decisive factor under distribution shift, such as transfer learning to new tissues, new species, or gene panels. Rather than identifying a single universally best tokenizer, Heimdall reveals that broad transferability depends on a small number of tokenization design axes that expose different biological priors to the model. These are precisely the conditions under which scFMs are most needed in practice, and our results reveal that current tokenization strategies introduce weaknesses that are not visible in standard benchmarks. We identify which components of tokenization drive generalization in each setting, show that some components from lower-ranked models actually outperform those from top-ranked ones, and demonstrate that hybrid tokenizers combining the best elements of existing designs can outperform any individual strategy. Together, these findings give both developers and users of scFMs a clearer picture of where current models are likely to succeed or fail, and a concrete path toward building more reliable ones through a principled framework for tokenizer design.

## Results

### Overview of H**eimdall**

In the Heimdall paradigm, the tokenizer maps a high-dimensional gene expression vector to a sequence of “tokens” as input for a sequence-based model such as a transformer. Heimdall distills tokenizers into three functional modules: *F*_**G**_, *F*_**E**_ and *F*_**C**_ (**Fig**. 1A). The gene embedding module *F*_**G**_ encodes the *identity* of a gene by incorporating prior biological knowledge; the expression embedding module *F*_**E**_ encodes the expression *value* of a gene by incorporating cell- or dataset-level statistics; and the aggregation module *F*_**C**_ integrates *F*_**G**_ and *F*_**E**_ outputs to assemble the final sequence-based representation of a cell. This formulation, which aggregates gene-level representations into a cell-level embedding, aligns with the standard formulation for universal representations of virtual cells [20]. For additional modularity, *F*_**C**_ is subdivided into order, sequence and reduce components (**Fig**. S1): order defines the intrinsic ordering of gene tokens, sequence selects which genes to include and constructs the token sequence, and reduce combines gene and expression encodings into a unified representation. Together, this modularization abstracts and generalizes key tokenization design decisions in the scFM literature. Additional details of the tokenizer abstraction can be found in **Methods**.

**Figure 1.**
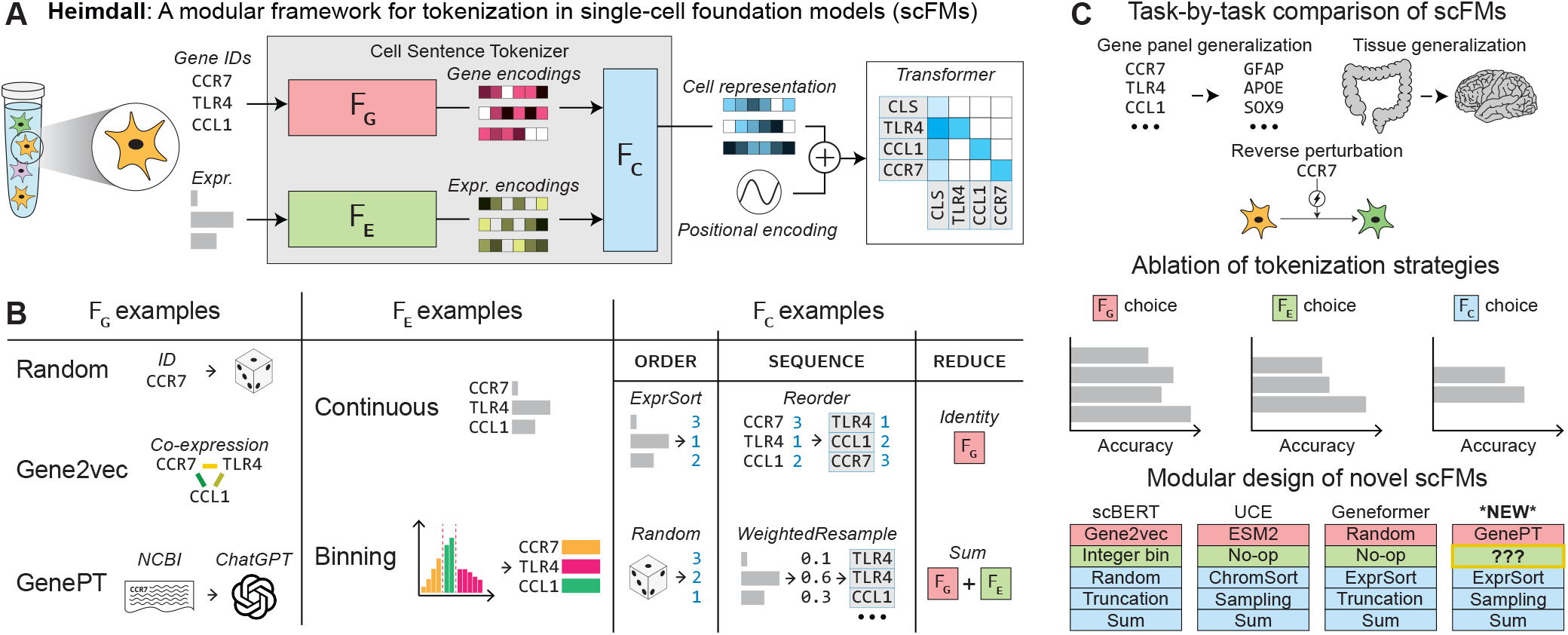
Overview of the Heimdall framework for single-cell foundation models (scFMs). **A**. Modular conceptualization of scFMs in Heimdall. Gene identities and expression values from each single-cell input are processed by a cell tokenization scheme (tokenizer) that generates a sequence-based “cell sentence”. The tokenizer is decomposed into three modules: a gene identity encoder (*F*_**G**_), an expression encoder (*F*_**E**_), and a cell constructor (*F*_**C**_). The tokenizer output is then passed to a sequence-based model (e.g., a transformer). **B**. Example implementations of the modular components, drawn from prior scFM literature. **C**. Applications of Heimdall for systematic design and evaluation of scFMs.

This interface enables Heimdall to reproduce input cell representation strategies from existing models. Using this abstraction, we reimplemented tokenizers from five top scFMs – scGPT [4], Gene-former [5], scFoundation [8], scBERT [9], and UCE [10] – denoted with a -tok suffix (e.g., Geneformertok) to distinguish them from their pretrained counterparts. These reimplementations yield concrete modular implementations of *F*_**G**_, *F*_**E**_ and *F*_**C**_ (**Fig**. 1B), with a comprehensive mapping of representative scFMs onto the Heimdall tokenizer decomposition summarized in **Table** S1.

The decoupling of tokenization from the sequence model and downstream tasks in Heimdall enables fair and measured comparisons of scFMs (**Fig**. 1C, top). By using a fixed set of hyperparameters for model training, Heimdall can control the effects of model and dataset size, model architecture, context length, positional encoding, and other elements. It also enables precise ablations of top-performing scFMs (**Fig**. 1C, middle) by swapping concrete implementations of tokenizer modules provided by Heimdall with the defaults used by the scFM. To attribute performance differences specifically to tokenization and enable scalable experiments, we forego pretraining and instead train all models from scratch. This allows us to compare tokenizers in a compute-efficient manner without making claims as to the pretraining objectives of existing scFMs. Beyond reproducing and dissecting existing scFMs, Heimdall facilitates modular design of novel tokenizers (**Fig**. 1C, bottom). Novel tokenizers can be constructed by mixing and matching Heimdall’s module implementations, while new modules can be seamlessly integrated into the framework.

We demonstrate Heimdall’s capabilities using three challenging single-cell transfer learning bench-marks – tissue shift (training on one organ, testing on another), cross species shift (training on species, testing on another), and gene-panel shift (predicting with unseen or sparsely overlapping genes). We further evaluate a paired-cell reverse perturbation prediction task, where the model infers a perturbation from a target cell state. Fundamentally, scFMs can only become useful and outperform bespoke methods once they learn robust, generalizable representations of cells. We consider these four downstream prediction tasks as interpretable, easy-to-optimize benchmarks that probe the cellular representations output by tokenizers to meaningfully capture and represent cellular identity.

Together, our development and applications highlight Heimdall’s potential to serve as a foundation for the rigorous, systematic, and measured development of high-performing scFMs.

### Evaluating the cross-tissue generalization capacity of scFMs

To evaluate how effectively different tokenizers support transfer learning across biological contexts, we constructed a cell type classification benchmark that measures cross-tissue generalization using a subset of the scTab CELLxGENE census [21] (referred to as the `sc-tab` dataset; see **Methods** for details). Models were trained on cells from the colon and small intestine and evaluated on cells from the brain across 15 shared cell types, providing a direct test of cross-tissue generalization (**Fig**. 2A).

**Figure 2.**
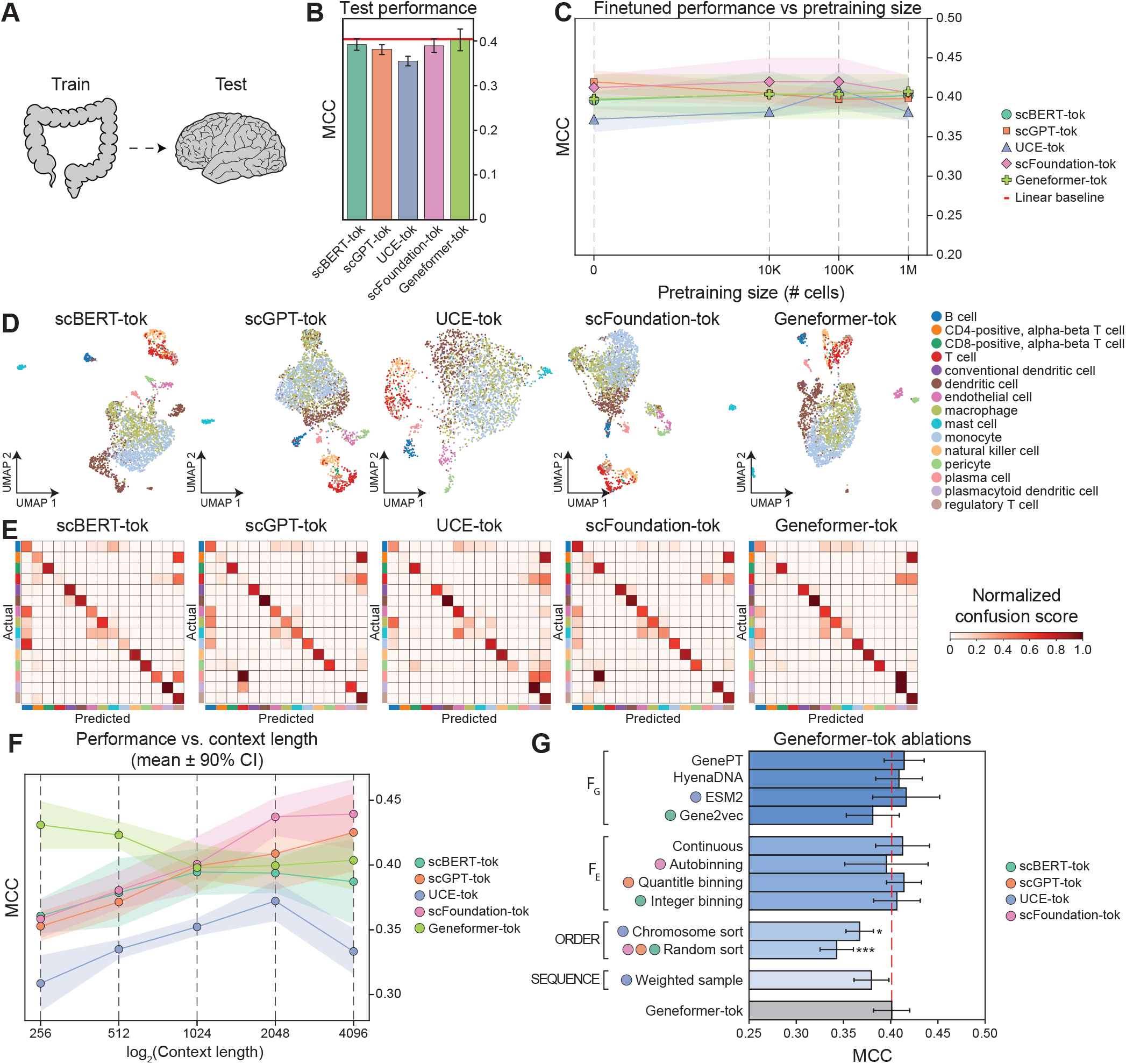
Comparison of scFM generalization performance on held-out tissue contexts using Heimdall. **A**. Schematic of the cross-tissue generalization task, where models are trained on colon and intestine cells and evaluated on brain cells. **B**. Performance (MCC; mean ± s.d. across *n* = 10 random seeds) of five scFMs tokenizers – scBERT-tok, scGPT-tok, UCE-tok, scFoundation-tok, and Geneformer-tok) – on the held-out brain dataset. **C**. Effect of pretraining scale: performance of tokenizers pretrained on progressively larger corpora and fine-tuned on `sc-tab` (x-axis: pretraining corpus size; 0 = no pretraining with random initialization). Error bars show s.d. across *n* = 5 random seeds. **D**. UMAP visualizations of the best-performing run for tokenization strategy, colored by ground-truth labels on the held-out brain dataset. **E**. Row-normalized confusion matrices of predicted versus true labels for each tokenizer. **F**. Performance across context lengths (256-4096 tokens; x-axis in log scale); error bars show s.d. across *n* = 5 random seeds. **G**. Module ablation of the best-performing tokenizer, Geneformer-tok, across Heimdall modules (*F*_**G**_, *F*_**E**_, order, sequence); error bars show s.d. across *n* = 6 random seeds. Significance was assessed using paired tests with Holm correction (p < 0.05, p < 0.01, p < 0.001).

We benchmarked tokenizers using matched conditions, including model size, context length, and no pretraining. We observed minimal performance variation across tokenizers, with Geneformer-tok performing best with a Matthews correlation coefficient (MCC) of 0.40, scBERT-tok and scFoundation-tok performing similarly (0.39), scGPT-tok slightly lower (0.38), and UCE-tok lowest (0.36). These differences were within run-to-run variability (s.d. 0.01-0.02) and overlapped with a linear baseline using raw gene expression (0.40), indicating that tokenizer choice has limited in-distribution impact for this task. Qualitative assessments using UMAP visualizations and confusion matrices corroborated this ranking (**Fig**. 2D-E).

To determine whether these conclusions extend to pretrained scFMs, we compared models trained from scratch to those pretrained on 10k, 100k, and 1M cells using a masked language modeling (MLM) objective and then finetuned (**Fig**. 2C). Across tokenizers, we observed limited benefit from MLM pretraining. scFoundation-tok showed small, monotonic gains with additional pretraining (±0.01), Geneformertok and scBERT-tok changed minimally, scGPT-tok declined modestly with larger pretraining sets, and UCE-tok improved modestly (+0.03). These results indicate that tokenizer choice – rather than pretraining – is the primary determinant of cross-tissue performance in this setting, with pretraining exerting only a marginal effect.

We next assessed how context length (tokens per cell) influences tokenizer performance in cross-tissue generalization (**Fig**. 2F). Performance increased with context length up to 2,048 tokens for scGPT-tok, scBERT-tok, and scFoundation-tok, after which it plateaued. This plateau can be attributed to the typical number of expressed genes per cell in `sc-tab`, with a median of 1,006 and an interquartile range of 598-1,637. Beyond 2,048 tokens, additional capacity primarily captures unexpressed or padded tokens, contributing limited new signal. Tokenization strategies that explicitly rank or resample genes exhibited distinctive scaling behavior: Geneformer-tok performance declined between 256 and 1,024 tokens before plateauing, whereas UCE-tok became unstable beyond 2,048 tokens.

To identify which tokenizer components drive cross-tissue generalization, we ablated the Geneformertok configuration within `sc-tab` using Heimdall (**Fig**. 2G). As all current tokenizers share the same reduce function, we varied the remaining modules (*F*_**G**_, *F*_**E**_, order, and sequence). Beyond the modules used in existing scFM tokenizers, we introduced three new variants: GenePT-based [6] and HyenaDNA-based gene identity embeddings for *F*_**G**_, and a continuous expression encoding for *F*_**E**_ (see **Methods**). Changes to *F*_**G**_ had limited effect: ESM2 (0.42 ± 0.03) and GenePT (0.41 ± 0.02) performed within the variability of the Geneformer-tok baseline with random embeddings (0.40 ± 0.02), indicating no meaningful *F*_**G**_-dependent differences for this task. Adding explicit expression encodings in *F*_**E**_ likewise did not yield consistent gains; scGPT-tok’s quantile binning matched the default No-Op *F*_**E**_ (0.41 ± 0.02 versus 0.40 ± 0.02). Within the order module, Geneformer-tok’s expression-based order remained best, showing significant gains over chromosome order (Holm-adjusted *p* < 0.05) and random order (*p* < 0.01) in paired tests across seeds; modifications to sequence did not surpass the default truncation-to-context strategy.

Overall, these ablations suggest that Geneformer-tok’s current formulation is already well specified for cross-tissue generalization, with limited room for further improvement via tokenization alone. Its advantage appears to arise primarily from the expression-sorting order, which implicitly injects expression information into the cell tokenization despite the absence of an explicit *F*_**E**_. Together, these results demonstrate how HEIMDALL facilitates systematic dissection of existing tokenizers and guides principled improvements.

### Evaluating the cross-species generalization capacity of scFMs

The transcriptional programs, regulatory modules, and signaling pathways that define cell identity are highly conserved across mammals [22]. However most scFMs are pretrained on a single species and inherit a species-specific gene vocabulary, making cross-species evaluation a stringent benchmark of tokenizer generalization. To test this, we trained models for each tokenizer on human cells and evaluated them on mouse cells without additional finetuning, using what we refer to as the `cross-species` dataset (**Fig**. 3A).

**Figure 3.**
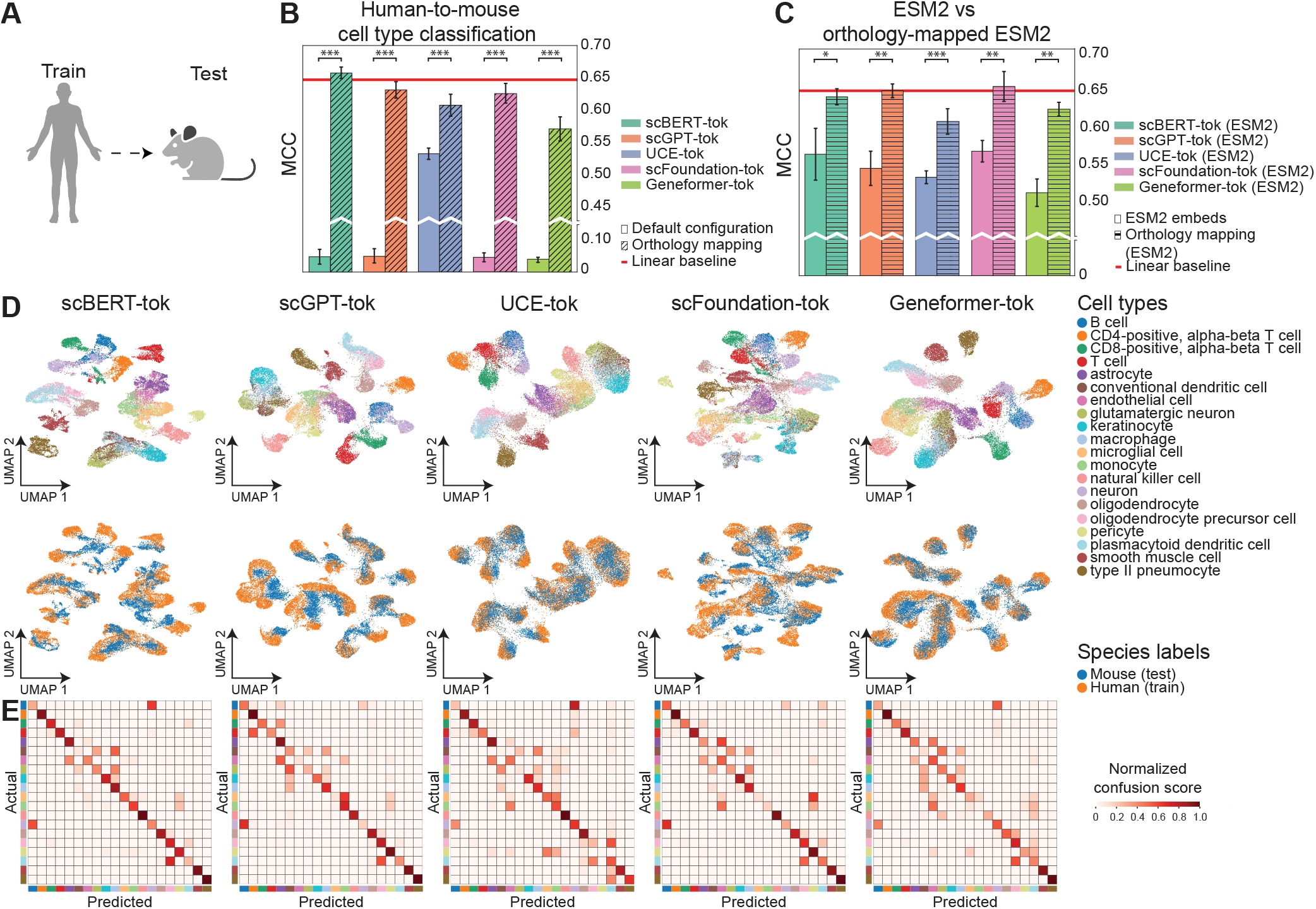
Evaluating the cross-species generalization capacity of scFMs. **A**. Schematic of the cross-species evaluation: models are trained on human cells and evaluated on mouse cells from matched cell types.**B**. Performance on the mouse test set for five scFM tokenizers, shown for each model’s default tokenization (solid) and after orthology-based human → mouse gene mapping (diagonal hatch). Bars show MCC across seeds (error bars: s.d., *n* = 5 random seeds); Significance was assessed using paired t-tests with Holm correction (p < 0.05, p < 0.01, p < 0.001). A red horizontal line marks the linear baseline. **C**. Ablations of the gene identity module (*F*_**G**_): replacing each model’s *F*_**G**_ with ESM2 embeddings (solid) or with ESM2 embeddings after orthology remapping (horizontal hatch). A red horizontal line marks the linear baseline. **D**. UMAP visualizations of the best-performing run for each tokenizer, colored by cell type (top) and species (bottom). **E**. Confusion matrices showing per-cell type precision and recall under cross-species transfer.

We evaluate two strategies for producing species-comparable cell representations. (1) *Sequence-based gene identity*: UCE-tok’s *F*_**G**_ represents each protein-coding gene as an ESM2 embedding of its protein sequence, yielding inherently species-agnostic representations. (2) *Orthology-based mapping*: for tokenizers lacking a sequence-based *F*_**G**_, mouse genes are mapped to their human orthologs prior to tokenization, aligning token identities with the human vocabulary used during training [23].

We first evaluated each tokenizer using its native configuration. scFoundation-tok, scGPT-tok, and Geneformer-tok all use randomly initialized embeddings for *F*_**G**_. scBERT-tok uses Gene2vec embeddings pretrained on human genes, with mouse-specific (out-of-vocabulary) genes represented by random vectors. In this no-mapping setting, only UCE-tok achieved informative cross-species performance, due to its sequence-derived *F*_**G**_ (**Fig**. 3B).

We next compared tokenizer performance while preserving each method’s *F*_**G**_, *F*_**E**_, and *F*_**C**_ modules under orthology-based mapping, which improved performance for all models (**Fig**. 3B). scBERT-tok performed best (0.66), followed by scGPT-tok (0.63), scFoundation-tok (0.63), UCE-tok (0.61), and Geneformer-tok (0.57). Notably, UCE-tok benefited from orthology mapping, even though it implicitly encodes orthologs. A linear baseline using the same orthology mapping performed similarly to the top-performing tokenizer scBERT-tok.

To directly test a species-agnostic *F*_**G**_, we performed an ablation in which each tokenizer’s original *F*_**G**_ was replaced with ESM2 embeddings, while *F*_**E**_ and *F*_**C**_ were left unchanged (UCE-tok already uses ESM2 and was not ablated). Under this shared setting, scFoundation-tok, scGPT-tok, scBERT-tok, and Geneformer-tok all outperformed UCE-tok (**Fig**. 3C), indicating that UCE’s cross-species advantage stems mostly from its sequence-derived *F*_**G**_, not from its *F*_**E**_ or *F*_**C**_.

Motivated by the gains from orthology mapping, we applied the same mapping to the ESM2 variants of each tokenizer. Across models, ESM2+orthology outperformed ESM2-only (**Fig**. 3C), consistent with improved alignment to the human training vocabulary and reduced out-of-vocabulary shift. Practically, orthology mapping and species-agnostic *F*_**G**_ are complementary: orthology mapping is advantageous where high-confidence maps exist, while a sequence-based *F*_**G**_ (e.g., UCE/ESM2) provides a robust, biologically grounded alternative when mappings are incomplete or uncertain.

UMAP visualizations of default scFM tokenizers under orthology mapping provide a qualitative view of each tokenizer’s embedding space, with cells colored by type and species (**Fig**. 3D). Cells of the same type within each species cluster tightly, and matched human-mouse cell types partially integrate in the embedding space. These qualitative patterns are reflected quantitatively in the confusion matrices (**Fig**. 3E).

Overall, UCE-tok emerged as the strongest mapping-free tokenizer for cross-species generalization, owing to its sequence-based *F*_**G**_ that confers species-comparable gene identities. When *F*_**G**_ is standardized (ESM2 across models) or orthology mapping is applied, tokenizers with stronger *F*_**E**_ and *F*_**C**_ components can surpass its performance. Nevertheless, for poorly annotated or non-model species lacking reliable orthology mappings, UCE-tok’s ESM2-based *F*_**G**_ remains the most practical and robust choice.

### Evaluating scFM generalization to spatial transcriptomics

Unlike scRNA-seq data, which typically measures most genes in the transcriptome, many imaging-based spatially-resolved transcriptomics (SRT) datasets measure only a small panel of genes (often fewer than 1,000), and cross-study gene panels are often disjoint or only partially overlapping [24, 25]. This panel mismatch creates a natural testbed for gene generalization, as datasets often share only a limited subset of genes (**Fig**. 4A).

**Figure 4.**
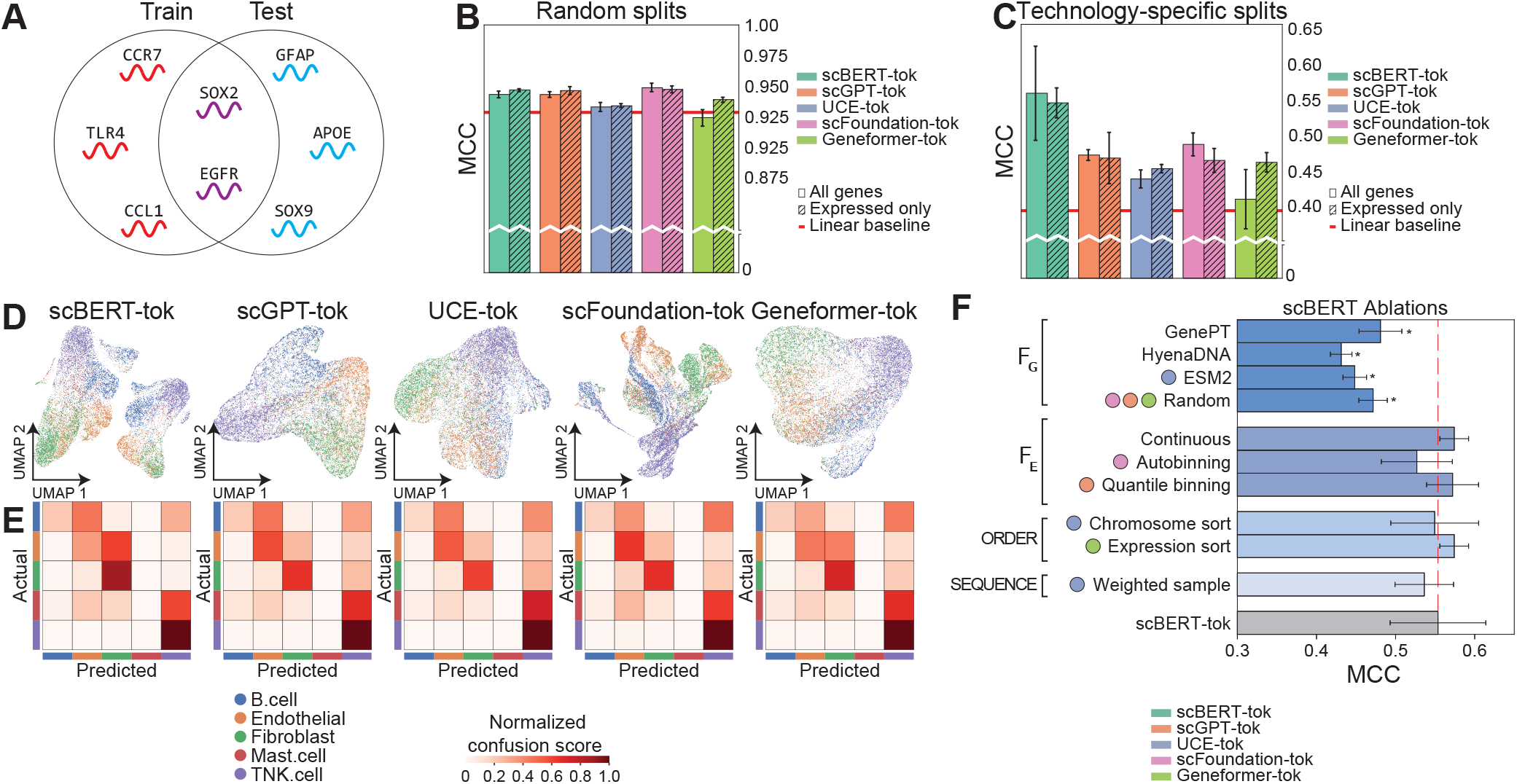
Spatial transcriptomics gene-panel generalization. **A**. Schematic of the gene-panel generalization task. **B**. Bar plots comparing tokenization strategies on random IID splits, reporting MCC for all genes and expressed-only genes. **C**. Bar plots comparing the same strategies under the `overlap-35` setting (35 shared genes). **D**. UMAP visualizations of the best-performing run for each scFM tokenizer. **E**. Confusion matrices for each tokenizer. **F**. Ablation of the best-performing tokenizer, scBERT-tok, across Heimdall modules. Significance was assessed using paired tests with Holm correction (p < 0.05, p < 0.01, p < 0.001). *Note:* In panels B-C, error bars show s.d. across 5 random seeds (*n* = 5); in panel F, error bars show s.d. across 6 random seeds (*n* = 6).

To first establish a within-dataset control before testing cross-panel generalization, we evaluated cell type classification on three ovarian cancer SRT samples with distinct gene panels [26] under independent and identically distributed (IID) splits, where training and test cells were drawn from the same distribution. Performance was uniformly high with minimal inter-model variance (**Fig**. 4B), indicating that tokenizer choice has little impact in IID settings.

We next evaluated performance under gene-panel shift using two non-IID benchmarks derived from three datasets: `overlap-35` (train panel 140 genes → test panel 280 genes; 35 shared) and `overlap-113` (train 280 genes → test 1,000 genes; 113 shared). On `overlap-35`, scBERT-tok performed best (0.55), followed by scGPT-tok, scFoundation-tok, and Geneformer-tok (0.47), with UCE-tok lowest (0.46) (**Fig**. 4C). All models exceeded a linear baseline (0.40), indicating that structured representations capture information beyond raw counts. UMAP visualizations and confusion matrices corroborated these rankings (**Fig**. 4D-E).

To account for tokenizer differences in handling non-expressed genes (see **Methods**), we evaluated both *expressed-only* and *all-genes* settings. This had little effect except for Geneformer-tok, which improved from 0.40 to 0.47 when restricted to expressed genes – consistent with its expression-ranking order being sensitive to many zeros in the input (**Fig**. 4C).

We next used Heimdall ‘s modularization to identify the source of scBERT-tok’s advantage. Because reduce is shared across tokenizers, we ablated only the remaining modules – *F*_**G**_, *F*_**E**_, and the *F*_**C**_ submodules order and sequence (**Fig**. 3F). *F*_**G**_ had the largest impact: Gene2vec remained strongest, with significant gains over GenePT, ESM2, and random (Holm-adjusted *p* < 0.05; paired tests across seeds). To probe this advantage, we analyzed the representational properties of each *F*_**G**_ (**Fig**. S3). Although Gene2vec embeddings showed weaker clustering of Gene Ontology terms than ESM2 or GenePT, co-expressed genes in the `overlap-35` benchmark had higher similarity in its embedding space (**Fig**. S3B-C). This pattern aligns with scBERT-tok’s performance margin, suggesting that co-expression priors stabilize representations for genes appearing only in the test set. Results on `overlap-113` mirrored these trends, with scBERT-tok leading and UCE-tok trailing (**Fig**. S2). Changing *F*_**E**_ yielded smaller but consistent gains – particularly continuous embeddings and quantile binning outperformed other *F*_**E**_ variants. Modifications to order, particularly the shift from random ordering to expression-based ordering, also led to small gains – while modifying sequence had minimal effect.

Together, these results show that under gene-panel shift, tokenizer design – particularly *F*_**G**_, with smaller gains from *F*_**E**_ and order– dominates performance. scBERT-tok consistently performs highest, UCE-tok lowest, and other tokenizers fall in between. These differences emerge only under generalization, not IID settings, indicating that a co-expression informed *F*_**G**_ and a well-designed *F*_**E**_ and order are critical for scFMs intended to generalize across gene panels.

### Single-cell perturbation inference via reverse perturbation

We next evaluated scFM tokenization strategies on the task of *reverse perturbation prediction*, which inverts the standard perturbation prediction formulation [4]. Instead of predicting a cell’s response to a given perturbation, the model is provided a target expression state and must identify perturbations likely to induce it. This formulation directly links cellular states to potential interventions, making it highly relevant for hypothesis generation and therapeutic target discovery.

We benchmarked tokenizers on the Norman single- and double-gene knockout dataset [27], which we processed into the `reverse-perturbation` dataset, framing the task as paired-cell classification where each instance consists of a control and its corresponding perturbed cell (**Fig**. 5A; task construction described in **Methods**). For paired-cell tasks, Heimdall introduces an AGGREGATOR module, which merges the individual embeddings of two cells provided by the transformer component into a single representation. For this task, we chose the asymmetric concatenation implementation of the AGGREGATOR module (see **Methods**). All models were benchmarked under matched conditions as in prior sections.

**Figure 5.**
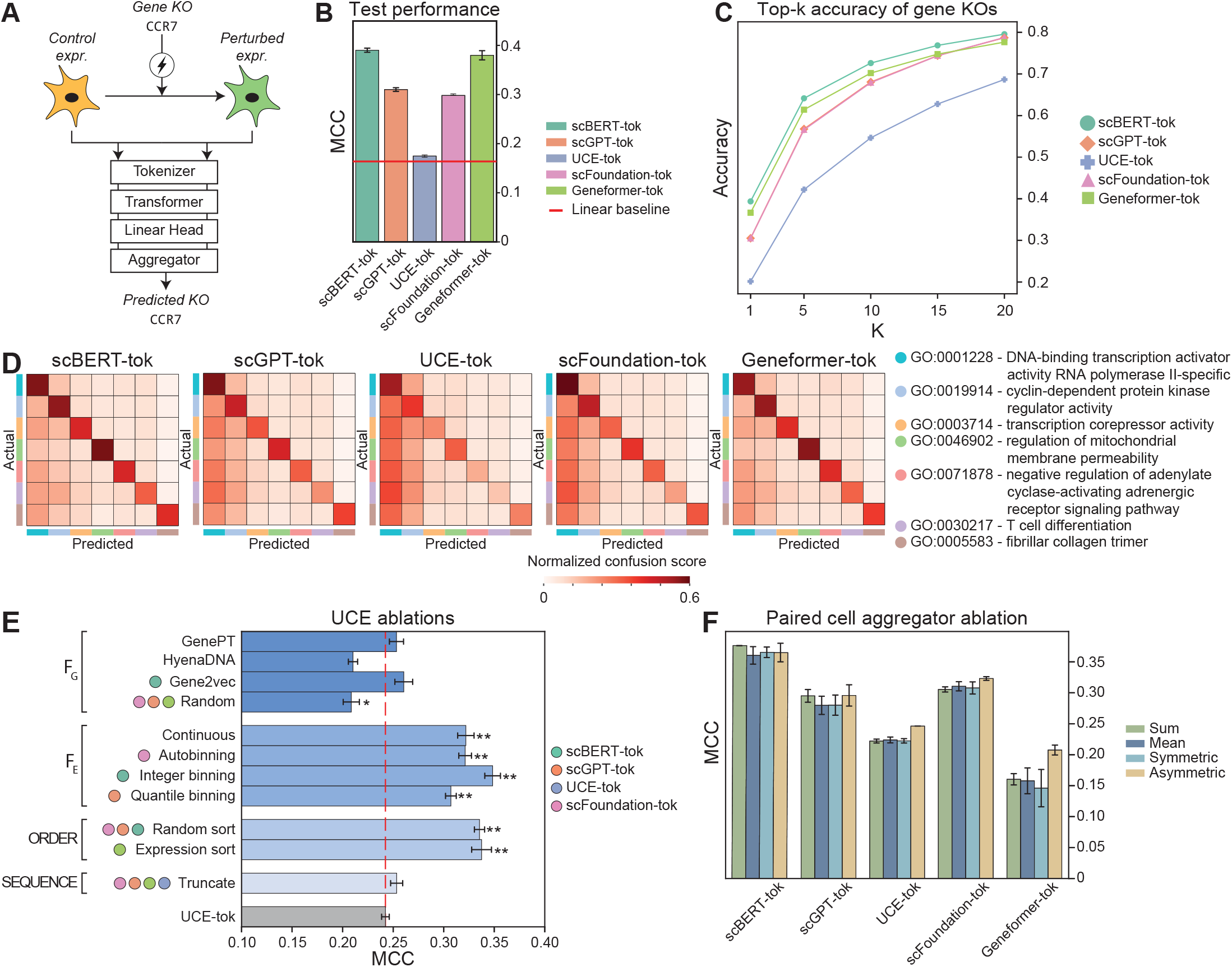
Comparison of performance on reverse perturbation prediction via Heimdall. **A**. Schematic of the reverse perturbation task, where perturbations are predicted from paired cells (control-perturbed pairs for single knockouts, or double knockouts where the first perturbation serves as the reference). **B**. Bar plots comparing cell tokenization strategies across scBERT, scGPT, UCE, scFoundation, and Geneformer, showing mean MCC ± s.d. across runs. **C**. Hit-rate (accuracy) of each method across k-thresholds. **D**. Gene-family heatmaps comparing predicted versus true knockout family memberships. **E**. Heimdall ablations of UCE-tok for each cell identity module. Significance was assessed using paired tests with Holm correction (p < 0.05, p < 0.01, p < 0.001). **F**. Ablation of paired-cell AGGREGATOR strategies across five scFMs. *Note:* Error bars show s.d. across *n* = 3 random seeds.

Performance varied substantially across tokenizers: scBERT-tok performed highest with a mean MCC of 0.39, followed by Geneformer-tok (0.38), scGPT-tok (0.31), scFoundation-tok (0.30), and UCE-tok (0.17) (**Fig**. 5B). All tokenizers outperformed a simple linear baseline that applied a difference-based AGGREGATOR to the paired expression profiles (0.16). We further quantified accuracy using hit-rate at threshold *k*, where *k* = 1 tests if the correct perturbation ranks first, and larger *k* values measure inclusion within the top-*k* predictions (**Fig**. 5C). All tokenizers showed a sharp increase in accuracy from *k* = 1 to *k* = 10, followed by smaller gains up to *k* = 20. Rankings were stable through *k* ≤ 15 (scBERT-tok highest, UCE-tok lowest), with scFoundation-tok slightly surpassing Geneformer-tok at *k* = 20.

Next, we evaluated whether each model’s predictions preserved gene-level semantics by comparing each model’s top-1 predicted knockout to the gene ontology (GO) family of the true gene [28] using row-normalized confusion matrices (**Fig**. 5D). scBERT-tok and Geneformer-tok showed the strongest concentration on the diagonal and fewest cross-family errors, suggesting that incorrect predictions often fall within the correct functional family. In contrast, scGPT-tok and scFoundation-tok distributed probability mass across adjacent families, while UCE-tok exhibited the most diffuse off-diagonal signal and biased predictions toward most prevalent families.

Using Heimdall’s per-module ablation, we examined why UCE-tok underperforms in reverse perturbation prediction (**Fig**. 5E). Modules lacking explicit expression information (*F*_**G**_ and sequence) had minor effects: within *F*_**G**_, ESM2 only exceeded random embeddings with significance (Holm-adjusted *p* < 0.05), while swaps to GenePT and Gene2vec did not. The largest improvements came from adding expression-aware modules (*F*_**E**_ and order), which follows naturally from UCE-tok’s default omission of expression encodings. In *F*_**E**_, continuous, autobinning, integer binning, and quantile binning each surpassed the UCE default (all Holm-adjusted *p* < 0.01), with integer binning (*F*_**E**_ from scBERT-tok) strongest (MCC of 0.35 versus 0.25). Further significant improvements came from replacing UCE-tok’s chromosome-based order with alternatives that better reflect expression (Geneformer-tok’s expression ordering) or by relaxing ordering assumptions with random ordering (Holm-adjusted *p* < 0.01). These results align with the original UCE study, which reported that adding chromosomal metadata led to a modest decrease in performance [10]. Finally, we performed an ablation of the paired-cell AGGREGATOR module, comparing (1) elementwise mean, (2) elementwise sum, (3) symmetric concatenation, and asymmetric concatenation (see **Methods**). For each tokenizer, asymmetric concatenation achieved either exceeded or matched the performance of other implementations, supporting its use in the main analyses (**Fig**. 5F).

Overall, rigorous evaluation with Heimdall reveals that changes to the *F*_**E**_ and order tokenization modules has the most impact on performance on the reverse perturbation task. For UCE-tok, which lacks expression encodings by default, including *any F*_**E**_ substantially improved performance. Combining scBERT-tok’s *F*_**E**_ and Geneformer-tok’s order yields the largest boost – highlighting these modules as key factors underlying top-model performance.

## Discussion

A central conclusion of this study is that tokenizer design is a key driver of transferability in scFMs, as it governs which biological priors are accessible to the model under distribution shift. Because single-cell data lack a canonical tokenization scheme, tokenizer design introduces inductive biases that strongly shape how well scFMs generalize across biological contexts. By decomposing scFM tokenizers into interpretable modules, Heimdall reveals that comparing whole models – as prior evaluations have done – can hide large differences in how well individual tokenizer choices actually perform. For example, although scBERT often ranks below other scFMs in earlier studies, Heimdall identifies that its Gene2vec-based gene identity encoding often outperforms those of other models in transfer learning settings. Together, this indicates that tokenizer design is an underappreciated source of fragility in current scFMs and a tractable target for developing more robust ones.

Our results further suggest that the capacity of scFMs for broad generalization is strongly influenced by a small number of tokenizer design axes. Across all settings, performance differences were driven primarily by gene identity, expression encoding, and ordering. This indicates that scFM behavior cannot be understood from architecture or scale alone – it also depends critically on *how* biological information is presented to the model. By making these design axes explicit, testable, and interchangeable, Heimdall provides a principled basis for designing tokenizers for specific deployment contexts and distribution shifts, rather than treating tokenization as an ad hoc modeling choice.

Several important directions remain for future work. Here, pretraining was examined only in a targeted ablation using a masked language modeling objective and a single transformer backbone. Future work could extend Heimdall to alternative pretraining objectives, larger corpora, other model architectures, and additional downstream tasks – including perturbation-response modeling, batch integration, gene regulatory network inference, and spatial tasks. More broadly, by making tokenizer design explicit and experimentally tractable, Heimdall enables systematic study of how tokenization interacts with model architecture, scale, and training corpus composition, all of which are important axes for advancing scFMs.

The importance of tokenizer design will only increase as scFMs move toward richer, multimodal representations. Looking ahead to *virtual cells* [20], transcriptomic profiles must be integrated with genomic, epigenomic, and proteomic analogs – and all of these signals must ultimately be translated into model inputs in a coherent, biologically meaningful way. Our use of HyenaDNA to encode gene identity illustrates one route to incorporate DNA sequence into tokenization, and deeper integration of regulatory and epigenomic signals may unlock the robust representations that virtual cells will require.

In summary, Heimdall identifies tokenization as a critical and underappreciated design axis in scFMs and provides a rigorous, reusable platform for developing the more robust scFMs that biomedical applications demand.

## Methods

### Tokenization paradigm for single-cell foundation models

To enable measured comparison and benchmarking of single-cell foundation models (scFMs), Heim-DALL represents their breadth and scope within a common functional paradigm. Using Heimdall, we reformulate several recently proposed scFMs under a unified, compositional framework.

For an input transcriptomic dataset 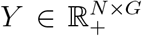 with gene features drawn from the set of all genes 𝒢_*all*_ and for a sequence embedding model with maximum sequence length *L* and hidden dimension *K*, the scFM function is defined as:

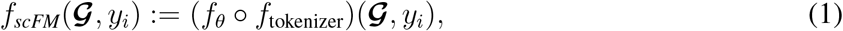

where *f*_tokenizer_ is the “cell tokenizer” that converts a cell’s gene expression vector *y*_*i*_ ∈ ℝ^*G*^ and gene set 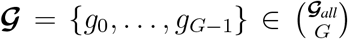 into an *L* × *K*-dimensional real-valued tokenization, and *f*_*θ*_ : ℝ ^*L*×*K*^ → ℝ ^*K*^ applies the transformer-based encoder to the tokenizer output. Here, *θ* are the encoder’s learnable parameters. The tokenizer is further decomposed as:

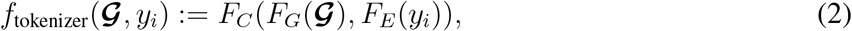

where 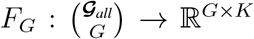 tokenizes gene identities, 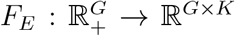 tokenizes gene expressions, and *F*_*C*_ : ℝ ^*G*×*K*^ × ℝ ^*G*×*K*^ → ℝ ^*L*×*K*^ combines the outputs of the *F*_*G*_ and *F*_*E*_.

To capture the breadth of pretraining tasks and downstream applications, we model task-specific decoders (“heads”) *f*_*ϕ*_ : ℝ ^*K*^ → ℒ, which map the scFM output to a label space ℒ. For example, for cell-type annotation over *C* cell types, the label space consists of one-hot *C*-dimensional vectors (i.e., ℒ = {0, 1}^*C*^ ); for gene expression reconstruction, 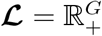.

**Fig**. S1 illustrates the Heimdall framework.

#### Abstraction of gene identity-based encodings via *F*_G_

The *F*_**G**_ component converts gene names/identifiers into *K*-dimensional real-valued embedding vectors, which are then forwarded to *F*_**C**_.

- **Random initialization**. Each gene is assigned a randomly initialized embedding vector, ensuring unique embeddings per gene.
- **Pretrained**. Several methods derive pretrained gene embeddings from external priors:
  - **ESM2**. The amino acid sequence of each gene’s protein product(s) is embedded with ESM2 [29]; multiple protein products are averaged.
  - **Gene2vec**. The Gene2vec [30] approach learns embeddings from gene coexpression patterns so co-expressed genes are nearby in embedding space.
  - **GenePT**. GenePT [6] uses the gene name and its NCBI text description as natural-language input to GPT-3.5 [31]; the output is taken as the embedding.
  - **HyenaDNA**. This approach embeds the gene’s DNA sequence (ENSEMBL release 112) with HyenaDNA [32]; the last token representation is used as the embedding.

If an embedding is unavailable for a given gene in a particular pretrained set, that gene is removed before subsequent tokenization steps. To resolve dimensionality mismatches between pretrained embeddings and the inherent dimensionality of the transformer model, we standardize and apply PCA to reduce all gene embeddings to 128 dimensions.

#### Abstraction of gene expression-based encodings via *F*_E_

The *F*_**E**_ component converts the expression values of a cell into a processed expression vector and a corresponding *K*-dimensional embedding vector; both are forwarded to *F*_**C**_.

- **No-op**. Outputs a constant zero vector per gene regardless of expression level (used by Geneformer and UCE).
- **Continuous**. A two-layer fully connected perceptron maps raw expression values to expression embeddings.
- **Quantile binning**. Expression values are assigned to *B* bins; each bin has a unique learnable embedding vector. Zero-valued expression is mapped to a dedicated “zero bin” (index 0). We use scGPT’s default *B* = 51.
- **Integer binning**. Expression values are partitioned into *B* bins by rounding each gene *y*_*ig*_ down and clipping to min (⌊*y*_*ig*_⌋, *B* − 1). We use scBERT’s default *B* = 7 for all benchmarks.
- **Autobinning**. Raw expression values are input to a two-layer fully connected perceptron that outputs *B* weights over learnable “autobin” embeddings; the final expression embedding is their weighted sum. We use scFoundation’s default *B* = 100 for all benchmarks.

Heimdall also allows the option of retaining or discarding zero-expression genes prior to *F*_**E**_, reflecting the scFM literature practices that include non-expressed genes in the input.

#### Abstraction of single-cell representation functions via the *F*_C_ module

The *F*_**C**_ module constructs the final tokenized single-cell representation by composing the outputs of *F*_**G**_ and *F*_**E**_. We partition *F*_**C**_ into three sequentially submodules:

1. The order function. Determines an intrinsic ordering of genes in the input cell, which sequence can use to place gene tokens into a meaningful sequence.
  - **Expression sorting**. Genes are ordered by their expression within the cell; expression values are normalized by the gene’s median across all cells prior to sorting (as in Geneformer).
  - **Chromosome sorting**. Genes are ordered by genomic coordinates in a two-stage process. First, genes are grouped into “chromosome blocks” by chromosome-of-origin; each block is delimited by start/end tokens, with start tokens unique to each chromosome and species. Because chromosomes have no intrinsic order, blocks are randomly permuted per cell. Second, within each block, genes are ordered by genomic start position. The permuted blocks are then concatenated to form the final sequence.
  - **Random ordering**. Genes receive a random permutation per cell.
2. The sequence function. This module constructs the sequence of tokens input to the final transformer architecture. It can use the gene order output by the order module to sequence genes. Beyond handling gene tokens, sequence may also inject metadata-based tokens to incorporate information complementary to the transcriptomic modality. For all sequence implementations, if the number of tokens exceeds the context length *L*, truncate to *L*; otherwise append `[PAD]` tokens to reach *L*.
  - **Truncation**. Place gene tokens according to order and truncate to context length *L*.
  - **Weighted sampling**. Sample genes with replacement, with probabilities proportional to their expression within the cell, until reaching *L*.
3. The reduce function. This module dictates how the *F*_**G**_ and *F*_**E**_ are combined for each gene token.
  - **Identity**. The *F*_**G**_ identity embeddings are directly used as the combined embeddings, and the *F*_**E**_ expression embeddings are ignored.
  - **Sum**. The *F*_**G**_ identity embeddings and *F*_**E**_ expression embeddings are summed element-wise to produce the combined embeddings.

Together, *F*_**G**_, *F*_**E**_ and *F*_**C**_ yield the overall tokenization scheme (tokenizer) for an scFM, applied individually to each cell in an scRNA-seq dataset.

#### Model architectures for processing tokenizer outputs

To process and extract information from a single-cell tokenization, scFMs use several machine learning model architectures. Most scFMs adopt a standard transformer architecture [33, 34], with stacked multihead attention layers applied to the gene-token sequence. Heimdall includes the following options:

- **Transformer**. The gene token sequence for each cell is input to a transformer with *n*_*layer*_ encoder blocks. Each block consists of a multi-head attention block followed by a fully connected multilayer perceptron.
- **Weighted token average**. For each cell, the weighted average of the *K*-dimensional token embeddings is used as the final embedding, with weights proportional to raw gene expression (as in GenePT [6] and CellPLM [35]).
- **Linear baseline**. The raw expression vector is fed directly to a single-layer fully-connected linear encoder to produce a *K*-dimensional cell embedding.

### Formulation of tasks

Heimdall flexibly supports the definition, configuration, and evaluation of diverse downstream tasks for scFMs. Defining a task requires specifying labels, a task-specific decoder (“head”) architecture, and a loss function (**Fig**. S1). Although not explored in this work, as a natural consequence of this abstraction, Heimdall natively supports multi-task learning.

#### Classification tasks

For classification tasks, precomputed class labels are provided for the input single-cell data, and the model predicts these labels for each cell. Examples include cell-type annotation (multiclass) and disease-risk prediction (binary).

- **Cell-type annotation**. A multi-class classification task in which the model predicts the cell type using the embedding of the `[CLS]` token. The loss is cross-entropy.

#### Regression tasks

For regression tasks, each cell is paired with one or more continuous targets (e.g., gene perturbation effect sizes, pseudotime, or per-gene expression), and the model predicts these values from the tokenized input. We formulate this as single- or multi-output regression and optimize mean squared error (MSE). Example applications include (i) perturbation prediction – estimating continuous gene knockout effects – and (ii) pretraining via expression reconstruction, where the model predicts gene expression and is trained with an MSE objective.

#### Pretraining tasks

Our framework supports pretraining tasks to train Heimdall on large datasets in a self-supervised manner. For such tasks, labels are generated during training by the dataset implementation within Heim-dall. Examples include:

- **Expression reconstruction**. Reconstruct the input gene expression vector *y*_*i*_ ∈ ℝ^*G*^ for each cell *i* (a regression task).
- **Masked language modeling**. As in NLP [36], masked language modeling (MLM) entails masking a random proportion of gene tokens in the cell sentence and replace them with `[MASK]` tokens. The model then predicts the identities of the masked tokens (a multiclass task).

#### Definition of paired-input tasks

Besides single-input tasks, Heimdall supports paired-input tasks in which each sample consists of a pair of cells. Each cell is passed through the model, and the resulting embeddings are reduced to a “pair embedding” using one of:

- **Mean**. Element-wise average.
- **Sum**. Element-wise sum.
- **Symmetric concatenation**. Concatenate embeddings and pass through a single-layer full connected perceptron to yield a *K*-dimensional output; repeat with the reverse order, then sum the two outputs to produce the pair embedding.
- **Asymmetric concatenation**. Concatenate embeddings in a fixed order and pass through a single-layer full-connected perceptron to yield a *K*-dimensional, order-sensitive output (no reserve-order pass).

### Extending Heimdall via novel implementations of tokenizer modules

Beyond reproducing existing scFM tokenizers, Heimdall provides standardized, well-documented interfaces for *F*_**G**_, *F*_**E**_, order, sequence, and reduce, enabling plug-and-play development of novel tokenization modules. We illustrate this extensibility by implementing and evaluating new modules for *F*_**G**_, *F*_**E**_, and reduce. First, we add a HyenaDNA-based *F*_**G**_ that provides sequence-derived gene-identity embeddings and include it in ablations throughout the paper. Second, we co-opt GenePT’s use of ChatGPT embeddings for NCBI text descriptions of genes as a novel *F*_**G**_– note that this contrasts from the original GenePT paper, which simply averages these gene tokens as opposed to using a transformer. Third, we introduce a continuous *F*_**E**_ that maps raw expression values to embeddings via a two-layer multilayer perceptron (MLP). While all five Heimdall-reproduced scFM tokenizers default to a sum reduce, tokenizers that omit explicit expression encodings in *F*_**E**_ (e.g., UCE and Geneformer) can also operate with an identity reduce. Comprehensive API documentation and examples are available in the Heimdall GitHub repository.

## Code Availability

Heimdall is released as an open-source Python package on the Python Package Index (`pip install sc-heimdall`). Furthermore, Heimdall is a fully open-source project hosted on GitHub: https://github.com/ma-compbio-lab/Heimdall

The GitHub repository includes reference implementations of the cell tokenizer components (*F*_**G**_, *F*_**E**_, order, sequence, reduce), model architectures, training/evaluation scripts, and the exact Hydra configuration files used in this study.

## Data Availability

- The `sc-tab` dataset [21] is publicly available for download at: https://github.com/theislab/scTab.
- The `overlap-113` and `overlap-35` [26] datasets are publicly available for download at: https://zenodo.org/records/12613839.
- The `cross-species` dataset was processed from CELLxGENE [12].
- The `reverse perturbation` [27] dataset is publicly available for download at:https://www.ncbi.nlm.nih.gov/geo/query/acc.cgi?acc=GSE133344.

A detailed description of how to download and construct these datasets can be found in the **Supplemental Information**.

## Acknowledgments

The authors would like to thank Yang Zhang, Junjie Tang, Wenduo Cheng, Shike Wang, Gün Kaynar and Xue Er Ding for helpful discussions and for assistance with dataset construction and curation. This work was supported, in part, by National Institutes of Health Common Fund 4D Nucleome Program grant UM1HG011593 (J.M.); National Institutes of Health Common Fund Cellular Senescence Network Program grant UH3CA268202 (J.M.); and National Institutes of Health grants R01HG007352 (J.M.), R01HG012303 (J.M.), R21DA061481 (J.M.), and R03OD039980 (J.M.). J.M. was additionally supported by the Ray and Stephanie Lane Professorship, a Guggenheim Fellowship from the John Simon Guggenheim Memorial Foundation, a Google Research Award, and a Single-Cell Biology Data Insights award from the Chan Zuckerberg Initiative. S.A. was supported by the NIH Training Grant T32EB009403 (2023-2024). N.H. was supported by a National Science Foundation Graduate Research Fellowship DGE2140739. S.K. and S.L. are Lane Fellows. The funders had no role in study design, data collection and analysis, decision to publish or preparation of the manuscript.

## Author Contributions

Conceptualization: E.H., S.A., N.H., S.K., and J.M.; Methodology: E.H., S.A., N.H., and S.K.; Software: E.H., S.A., N.H., and S.K.; Investigation: E.H., S.A., N.H., R.L., E.T., S.L., M.Y., S.K., and J.M.; Writing: E.H., S.A., N.H., S.K., and J.M.; Funding Acquisition: J.M.

## References

[1] Jumper, J. et al. Highly accurate protein structure prediction with alphafold. Nature 596, 583–589 (2021).

[2] Dalla-Torre, H. et al. Nucleotide transformer: building and evaluating robust foundation models for human genomics. Nature Methods 1–11 (2024).

[3] Moor, M. et al. Foundation models for generalist medical artificial intelligence. Nature 616, 259– 265 (2023).

[4] Cui, H. et al. scGPT: toward building a foundation model for single-cell multi-omics using generative AI. Nature Methods 21, 1470–1480 (2024).

[5] Theodoris, C. V. et al. Transfer learning enables predictions in network biology. Nature 618, 616–624 (2023).

[6] Chen, Y. & Zou, J. Simple and effective embedding model for single-cell biology built from chatgpt. Nature Biomedical Engineering 9, 483–493 (2025).

[7] Levine, D. et al. Cell2sentence: teaching large language models the language of biology. BioRxiv 2023–09 (2024).

[8] Hao, M. et al. Large-scale foundation model on single-cell transcriptomics. Nature Methods 21, 1481–1491 (2024).

[9] Yang, F. et al. scBERT as a large-scale pretrained deep language model for cell type annotation of single-cell rna-seq data. Nature Machine Intelligence 4, 852–866 (2022).

[10] Rosen, Y. et al. Universal cell embeddings: A foundation model for cell biology. bioRxiv 2023–11 (2023).

[11] Tarhan, L. et al. Single Cell Portal: an interactive home for single-cell genomics data. BioRxiv (2023).

[12] Program, C. C. S. et al. CZ CELLxGENE Discover: a single-cell data platform for scalable exploration, analysis and modeling of aggregated data. Nucleic Acids Research 53, D886–D900 (2025).

[13] Regev, A. et al. The human cell atlas. eLife 6, e27041 (2017).

[14] Liu, T., Li, K., Wang, Y., Li, H. & Zhao, H. Evaluating the utilities of foundation models in single-cell data analysis. bioRxiv 2023–09 (2023).

[15] Ahlmann-Eltze, C., Huber, W. & Anders, S. Deep-learning-based gene perturbation effect prediction does not yet outperform simple linear baselines. Nature Methods 22, 1657–1661 (2025).

[16] Bendidi, I. et al. Benchmarking transcriptomics foundation models for perturbation analysis : one PCA still rules them all. arXiv (2024).

[17] Boiarsky, R. et al. Deeper evaluation of a single-cell foundation model. Nature Machine Intelligence 6, 1443–1446 (2024).

[18] Kedzierska, K. Z., Crawford, L., Amini, A. P. & Lu, A. X. Zero-shot evaluation reveals limitations of single-cell foundation models. Genome Biology 26, 101 (2025).

[19] Lotfollahi, M. Toward learning a foundational representation of cells and genes. Nature Methods 21, 1416–1417 (2024).

[20] Bunne, C. et al. How to build the virtual cell with artificial intelligence: Priorities and opportunities. Cell 187, 7045–7063 (2024).

[21] Fischer, F. et al. scTab: Scaling cross-tissue single-cell annotation models. Nature Communications 15, 6611 (2024).

[22] Breschi, A., Gingeras, T. R. & Guigó, R. Comparative transcriptomics in human and mouse. Nature Reviews Genetics 18, 425–440 (2017).

[23] stefanpeidli/mousipy: Translates an AnnData object with mouse gene symbols into one with human gene symbols by mapping orthologs from biomart. URL https://github.com/stefanpeidli/mousipy.

[24] Hu, Y. et al. Benchmarking clustering, alignment, and integration methods for spatial transcriptomics. Genome Biology 25, 212 (2024).

[25] Haber, E., Deshpande, A., Ma, J. & Krieger, S. Unified integration of spatial transcriptomics across platforms. In International Conference on Research in Computational Molecular Biology, 371–375 (Springer, 2025).

[26] Yeh, C. Y. et al. Mapping spatial organization and genetic cell-state regulators to target immune evasion in ovarian cancer. Nature Immunology 25, 1943–1958 (2024).

[27] Norman, T. M. et al. Exploring genetic interaction manifolds constructed from rich single-cell phenotypes. Science 365, 786–793 (2019).

[28] Klopfenstein, D. et al. GOATOOLS: A python library for Gene Ontology analyses. Scientific Reports 8, 1–17 (2018).

[29] Lin, Z. et al. Evolutionary-scale prediction of atomic-level protein structure with a language model. Science 379, 1123–1130 (2023).

[30] Du, J. et al. Gene2vec: distributed representation of genes based on co-expression. BMC Genomics 20, 82 (2019).

[31] Shen, H. et al. Generative pretraining from large-scale transcriptomes for single-cell deciphering. iScience 26 (2023).

[32] Nguyen, E. et al. Hyenadna: Long-range genomic sequence modeling at single nucleotide resolution. Advances in Neural Information Processing Systems 36, 43177–43201 (2023).

[33] Vaswani, A. et al. Attention is all you need. Advances in neural information processing systems 30 (2017).

[34] Szałata, A. et al. Transformers in single-cell omics: a review and new perspectives. Nature Methods 21, 1430–1443 (2024).

[35] Wen, H. et al. CellPLM: Pre-training of cell language model beyond single cells. In The Twelfth International Conference on Learning Representations (2024).

[36] Devlin, J., Chang, M.-W., Lee, K. & Toutanova, K. BERT: Pre-training of deep bidirectional transformers for language understanding. In Proceedings of the 2019 Conference of the North American Chapter of the Association for Computational Linguistics: Human Language Technologies (Vol. 1), 4171–4186 (2019).

