## Supplemental Information for "HEIMDALL: Disentangling tokenizer design for robust transfer in single-cell foundation models"

#### A Supplementary Results

##### A.1 Explaining the influence of gene identity embeddings on scFM performance

To probe why some gene identity encoders ($F_{\mathbf{G}}$) outperform others, we first visualized each $F_{\mathbf{G}}$ using UMAP, colored by Gene Ontology (GO) terms (**Fig.** S3A). GenePT and ESM2 embeddings showed clearer GO-related structure than Gene2vec or HyenaDNA.

We quantified these patterns with two metrics. First, a GO-term cohesion score tested whether genes sharing a GO term cluster more tightly than size-matched random sets (**Fig.** S3B, left; **Methods**). Second, we applied Louvain clustering and computed the adjusted Rand index (ARI) between clusters and GO terms (**Fig.** S3B, right).

Although ESM2 exhibits strong GO term cohesion, it performs poorly on the `overlap-35` dataset. Conversely, Gene2vec shows low cohesion and purity yet remains one of the top-performing $F_{\mathbf{G}}$s. Thus, alignment with GO alone does not explain $F_{\mathbf{G}}$ performance as a cell tokenization strategy, despite GO reflecting functional similarity.

We next asked whether gene co-expression better accounts for $F_{\mathbf{G}}$ performance by comparing pairwise similarities in embedding space to co-expression in `overlap-35` and `sc-tab` (**Fig.** S3C). These correlation-based metrics aligned more closely with $F_{\mathbf{G}}$ rankings, with Gene2vec and GenePT scoring highly. Notably, random embeddings scored poorly here yet still outperformed ESM2 and HyenaDNA in cell type classification, indicating that correlation to co-expression is not the sole determinant of downstream performance.

While $F_{\mathbf{G}}$ rankings by co-expression were broadly consistent across datasets, Gene2vec scored markedly higher on `overlap-35`, mirroring its superior classification accuracy, whereas GenePT was more competitive on `sc-tab`. These results suggest that $F_{\mathbf{G}}$s capturing co-expression relationships are better positioned to compensate for technical dropout, improving cell type classification.

In summary, gene encoders that capture co-expression patterns, rather than functional similarity alone, are better aligned with downstream performance, underscoring the importance of data-driven encoding properties for cell-level tasks.

### B Supplementary Methods

#### B.1 Benchmark and analysis details

We provide details of how each dataset was constructed and analyzed, including preprocessing, train, validation, and test splits, and any other dataset-specific attributes.

##### *Details of the `sc-tab` benchmark and analysis*

The `sc-tab` dataset was constructed following the same preprocessing and filtering procedures described in scTab [1]. Briefly, we downloaded data from CELLxGENE [2] and filtered to include only primary data generated using 10x Genomics protocols with at least 5,000 cells per cell type. We then selected the most common cell types from tissues with the largest sample sizes. To evaluate cross-tissue generalization, we used gastrointestinal tissues for training and brain tissues for validation and testing (train $n = 36900$; val $n = 2769$; test $n = 2759$).

##### *Details of the `cross-species` benchmark and analysis*

The `cross-species` dataset was also downloaded from CELLxGENE. We first applied a preprocessing pipeline similar to the `sc-tab` dataset: retaining primary data generated by 10x Genomics protocols and requiring at least 5,000 cells per cell type. From these, we selected 20 of the most common and diverse human cell types, then queried CELLxGENE for mouse cells corresponding to the same 20 categories and subsampled 1,000 cells per type, yielding a final dataset of 40,000 cells. Human cells were used for training, while mouse cells were randomly split into validation and test sets.

##### *Details of the `overlap-113` and `overlap-35` benchmark and analysis*

The `overlap-35` and `overlap-113` datasets are derived from a spatial transcriptomics dataset of human ovarian cancer [3]. In the original study, data were assayed using five different gene panels. We use a subset originally labeled "Validation 1", "Validation 2", and "Test 2", where all cells share the same anatomical region (adnexa), survival outcome, cancer stage, and diagnostic age. "Validation 1" was captured using Xenium (240 genes), "Validation 2" with MERFISH (140 genes), and "Test 2" with CosMx (1000 genes). We did not account for other confounding factors such as treatment or tumor genetics. Original cell type labels were assigned to each dataset separately using a common procedure and unified set of cell type labels.

The `overlap-35` dataset uses "Validation 1" for training and "Validation 2" for validation/testing (35 shared genes). The `overlap-113` dataset uses "Validation 2" for training and "Test 2" for validation/testing (113 shared genes). For each original datasets, we randomly subsampled 10,000 cells per each cell type shared between the train and the test splits; for Mast cells (fewer than 10,000 per dataset), we used all available cells.

##### *Details of the* `reverse-perturbation` *benchmark and analysis*

To evaluate the model's ability to predict the effects of genetic perturbations, we curated a task based on the Perturb-seq dataset from Norman et al. [4] This dataset contains scRNA-seq profiles of K562 cells with 287 single and dual-gene knockouts (KOs) mediated by CRISPR-Cas9, as well as non-targeting control cells. We formulated the task as predicting the differential effect of a gene KO relative to a defined baseline. For single-gene KOs (e.g., gene A), we created pairs of perturbed cells (A-KO) and control cells (NegCtrl). For dual-gene KOs, we model double knockouts as conditional on a single knockout; that is, for a dual-gene KO (A, B), we create pairs by treating A-KO (or, without loss of generality, B-KO) as the control cells and (A+B)-KO as the perturbed cells . To avoid leakage, we first partitioned cells within each unique perturbation condition into training (60%), validation (20%), and test (20%) sets. Then, for each differential task, we randomly sampled a fixed number of pairs (8,000 train; 1,000 validation, and 1,000 test).

We also created a compact Norman subset for reverse-perturbation ablation. We enumerated KO "conditions" in the train/val/test matrices – single KOs ("A") and double KOs ("A|B") – and greedily selected $k = 20$ genes maximizing weighted coverage of observed conditions across splits. We retained all cells from selected single-gene KOs and from double-gene KOs whose partners were both in-panel, and kept non-targeting controls (NegCtrl). We rebuilt a compact KO index (NegCtrl $\to 0$, in-panel KOs $\to 1..K$) and excluded NegCtrl rows from supervision so controls serve only as baselines. Finally, we defined a boolean mask identifying all paired examples included in train/val/test and verified that only non-control KOs contribute to supervised labels.

To assess correspondence between predicted and true gene KOs, we grouped genes into functional "families". Briefly, we mapped gene symbols → Entrez [5]; loaded the human GO DAG and Gene2GO; pruned each gene's annotations to specific BP/MF/CC terms (depth/IC-filtered, leaves only); kept top-IC terms per namespace; computed pairwise semantic similarity using Lin's metric with namespace weighting and an overlap scalar; and clustered the matrix to define families, labeled via GO enrichment. For the figure, we mapped true and predicted genes to these families and summarized correspondence using Top-1 majority-family agreements and row-normalized family confusion matrices.

##### *Statistical testing*

Statistical testing was used to assess the significance of per-module ablations in HEIMDALL. Significance was assessed by comparing baseline tokenizers to module ablations across downstream tasks using paired tests across random seeds. We tested normality of per-seed differences with the Shapiro-Wilk test; if normality was not rejected ($p \geq 0.05$), we used a two-sided paired t-test; otherwise, a two-sides Wilcoxon signed-rank test. P-values were adjusted for multiple comparisons using Holm's method within each ablation family ($F_{\mathbf{G}}$, $F_{\mathbf{E}}$, ORDER, SEQUENCE) with $\alpha = 0.05$. Bars show mean across seeds; error bars denote standard deviation. Stars denote adjusted significance: $^{*}p < 0.05,^{**} p < 0.01,^{***} p < 0.001$.

##### B.2 Configuration of HEIMDALL for benchmarking and ablation experiments

Our framework uses Hydra [6] to configure all aspects of a HEIMDALL experiment, including the tokenizer function, model architecture, task, and all other hyperparameters. Each configuration inherits from modular YAML files in configs/, allowing composable overrides. Unless otherwise stated, all experiments were AdamW with a cosine-annealing learning-rate schedule. The model is a Transformer with two encoder layers and four attention heads per layer, with FlashAttention enabled. For each benchmark, we performed a limited hyperparameter sweep over learning rate and batch size to ensure stability across datasets, testing learning rates $1 \times 10^{-2}, 1 \times 10^{-3}, 3 \times 10^{-4}$ and batch sizes in $32, 64$, selecting the configuration with the best validation Matthews correlation coefficient (MCC).

*`sc-tab`*

Configuration file: `/configs/experiments/sctab_split1_all.yaml`

- **Task**: cell-type classification `/configs/tasks/new_sctab_split.yaml`
  - **Objective**: multiclass classification
  - **Metrics**: MCC
  - **Split strategy**: cross-tissue transfer (train: colon + intestine; test: brain)
  - **Type**: single-instance dataset

*`cross-species`*

Configuration file: `/configs/experiments/sctab_split1_all.yaml`

- **Task**: cell-type classification `/configs/tasks/new_sctab_split.yaml`
  - **Objective**: multiclass classification
  - **Metrics**: MCC
  - **Split strategy**: cross-species transfer (train: human; test: mouse)
  - **Type**: single-instance dataset

*`overlap-35`*

Configuration file: `/configs/experiments/spatial_cancer_split1.yaml`

- **Task**: cell-type classification `/configs/tasks/spatial_cancer_split.yaml`
  - **Objective**: multiclass classification
  - **Metrics**: MCC
  - **Split strategy**: cross-gene panel transfer (train: Validation 1, Xenium, 240 genes; test: Validation 2, MERFISH, 140 genes; 35 shared genes)
  - **Type**: single-instance dataset

*`overlap-113`*

Configuration file: `/configs/experiments/spatial_cancer_split2.yaml`

- **Task**: cell-type classification `/configs/tasks/spatial_cancer_split.yaml`
  - **Objective**: multiclass classification
  - **Metrics**: MCC
  - **Split strategy**: cross-gene panel transfer (train: Validation 2, MERFISH, 140 genes; test: Test 2, CosMx, 1000 genes; 113 shared genes)
  - **Type**: single-instance dataset

*`reverse-perturbation`*

Configuration file: `/configs/experiments/reverse_perturbation.yaml`

- **Task**: cell-type classification `/configs/tasks/reverse_perturbation.yaml`
  - **Objective**: multiclass classification
  - **Metrics**: MCC
  - **Split strategy**: cross-gene panel transfer (train: Validation 2, MERFISH, 140 genes; test: Test 2, CosMx, 1000 genes; 113 shared genes)
  - **Type**: paired-instance dataset

#### Supplementary Figures

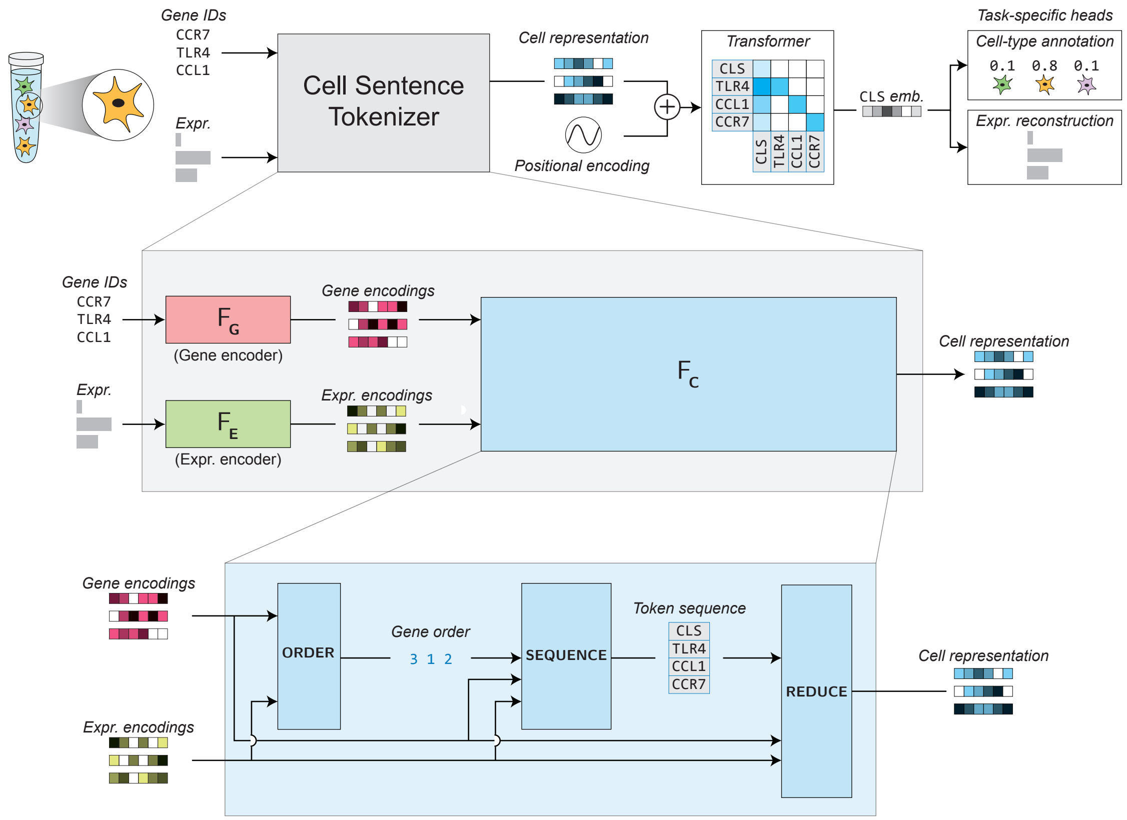


**Figure S1: Extended overview of the HEIMDALL framework.** Modular conceptualization of the cell tokenizer, decomposing the $F_{\mathrm{C}}$ module into its subcomponents: ORDER, SEQUENCE, and REDUCE. Gene identity encodings from $F_{\mathrm{G}}$ and expression encodings from $F_{\mathrm{E}}$ serve as inputs to the cell constructor $F_{\mathrm{C}}$. Within $F_{\mathrm{C}}$, ORDER first determines the gene ordering; the resulting order and expression encodings are then passed to SEQUENCE, which selects the subset of genes to include in the sequence. Finally, the ordered token sequence of gene and expression embeddings is passed to REDUCE, which integrates them into a unified cell level representation.

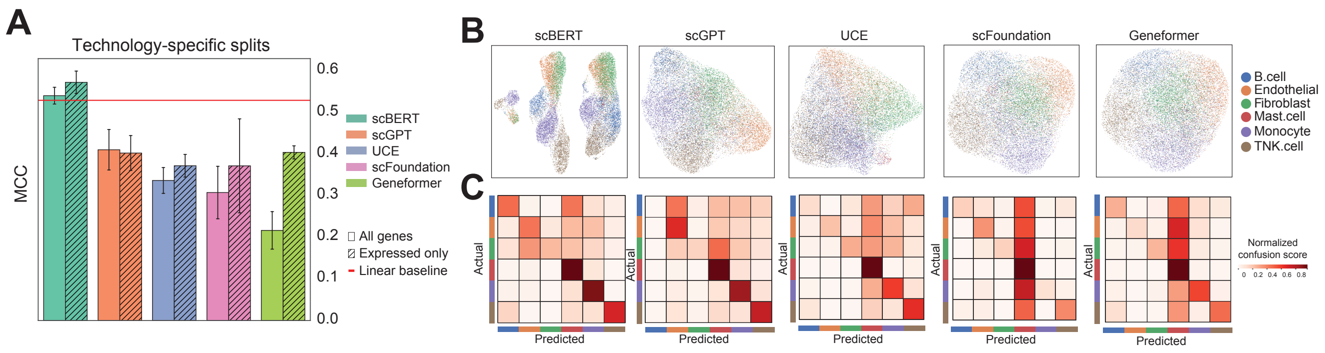


**Figure S2: Spatial transcriptomics gene-panel generalization A.** Bar plots comparing tokenization strategies across five scFMs (scBERT, scGPT, UCE, scFoundation, and Geneformer) on the `overlap-113` cross-technology split, reporting performance for all genes vs. expressed genes across $n = 5$ random seeds. **B.** UMAP visualizations of the best-performing run for each scFM tokenization strategy. **C.** Confusion matrices for each tokenizer.

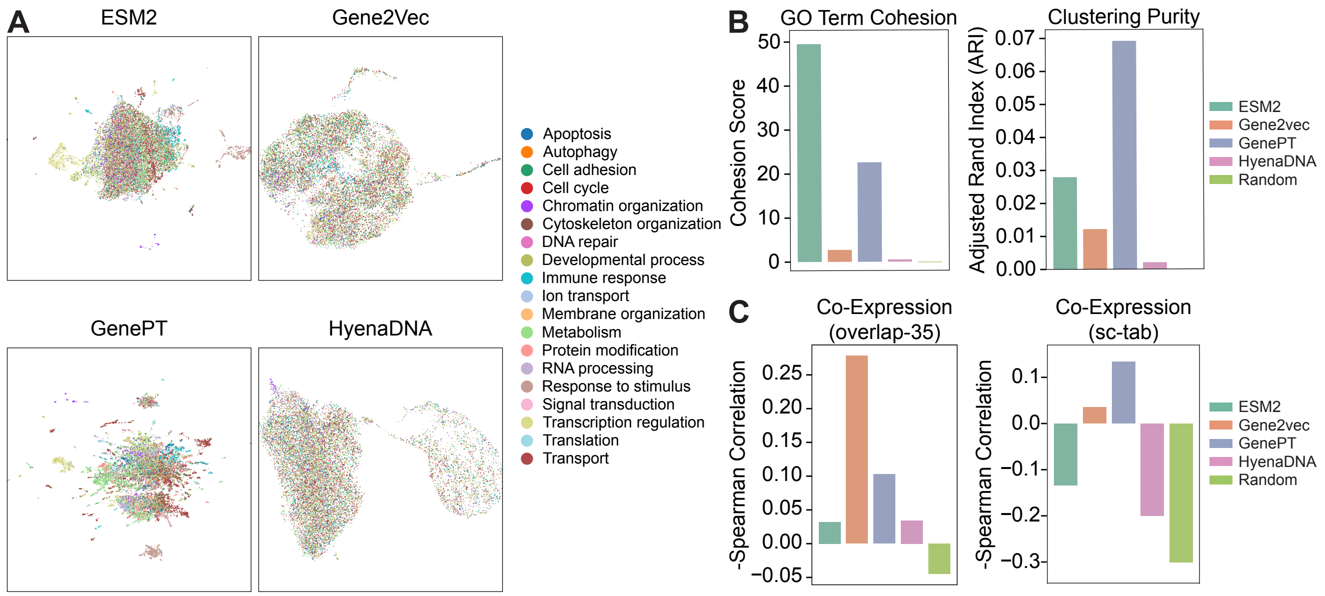


**Figure S3: Comparison of gene-encoding strategies ($F_G$). A.** UMAP projection of gene embeddings from four encoding strategies (excluding Random), after reducing all embeddings to a common dimensionality with PCA. **B.** Bar plots summarizing GO-term grouping performance for each $F_G$. Left: cohesion score, defined as the negated z-score of within-term gene distances relative to random sets of the same size. Right: adjusted Rand Index (ARI) obtained by applying Louvain clustering to the embeddings and comparing clusters to GO annotations. **C.** Bar plots showing how well pairwise distances in each embedding space recapitulate experimental relationships: two plots for the `overlap-35` dataset (gene-to-gene coexpression and spatial correlation) and one for the `sc-tab` dataset (expression correlation). Each reports the Spearman correlation between embedding distance and the corresponding experimental metric across all five $F_G$s.

### Supplementary Tables

**Table S1:** Comparison of tokenizers for single-cell foundation models' evaluated in this work.

| **Model** | $F_{\mathbf{G}}$ | $F_{\mathbf{E}}$ | $F_{\mathbf{O}}$ | $F_{\mathbf{S}}$ | $F_{\mathbf{R}}$ |
|---|---|---|---|---|---|
| scBERT [7] | Gene2vec [8] | Integer bin | Random | Truncation | Sum |
| scGPT [9] | Random | Quantile bin | Random | Truncation | Sum |
| UCE [10] | ESM2 [11] | No-Op | Chromosome | Sampling | Sum |
| scFoundation [12] | Random | Autobin | Random | Truncation | Sum |
| Geneformer [13] | Random | No-Op | Expression | Truncation | Sum |

### References


[1] Fischer, F. *et al.* scTab: Scaling cross-tissue single-cell annotation models. *Nature Communications* **15**, 6611 (2024).

[2] Program, C. C. S. *et al.* CZ CELLxGENE Discover: a single-cell data platform for scalable exploration, analysis and modeling of aggregated data. *Nucleic Acids Research* **53**, D886–D900 (2025).

[3] Yeh, C. Y. *et al.* Mapping spatial organization and genetic cell-state regulators to target immune evasion in ovarian cancer. *Nature Immunology* **25**, 1943–1958 (2024).

[4] Norman, T. M. *et al.* Exploring genetic interaction manifolds constructed from rich single-cell phenotypes. *Science* **365**, 786–793 (2019).

[5] Schuler, G. D., Epstein, J. A., Ohkawa, H. & Kans, J. A. Entrez: Molecular biology database and retrieval system. In *Methods in Enzymology*, vol. 266, 141–162 (Elsevier, 1996).

[6] Yadan, O. Hydra - a framework for elegantly configuring complex applications. GitHub (2019). URL https://github.com/facebookresearch/hydra.

[7] Yang, F. *et al.* scBERT as a large-scale pretrained deep language model for cell type annotation of single-cell rna-seq data. *Nature Machine Intelligence* **4**, 852–866 (2022).

[8] Du, J. *et al.* Gene2vec: distributed representation of genes based on co-expression. *BMC Genomics* **20**, 82 (2019).

[9] Cui, H. *et al.* scGPT: toward building a foundation model for single-cell multi-omics using generative AI. *Nature Methods* **21**, 1470–1480 (2024).

[10] Rosen, Y. *et al.* Universal cell embeddings: A foundation model for cell biology. *bioRxiv* 2023–11 (2023).

[11] Lin, Z. *et al.* Evolutionary-scale prediction of atomic-level protein structure with a language model. *Science* **379**, 1123–1130 (2023).

[12] Hao, M. *et al.* Large-scale foundation model on single-cell transcriptomics. *Nature Methods* **21**, 1481–1491 (2024).

[13] Theodoris, C. V. *et al.* Transfer learning enables predictions in network biology. *Nature* **618**, 616–624 (2023).
